# Multimerization of *Homo sapiens* TRPA1 ion channel cytoplasmic domains

**DOI:** 10.1101/466060

**Authors:** Gilbert Q. Martinez, Sharona E. Gordon

**Affiliations:** Department of Physiology and Biophysics, University of Washington, Seattle, Washington, United States of America

## Abstract

The transient receptor potential Ankyrin-1 (TRPA1) ion channel is modulated by myriad noxious stimuli that interact with multiple regions of the channel, including cysteine-reactive natural extracts from onion and garlic which modify residues in the cytoplasmic domains. The way in which TRPA1 cytoplasmic domain modification is coupled to opening of the ion-conducting pore has yet to be elucidated. The cryo-EM structure of TRPA1 revealed a tetrameric C-terminal coiled-coil surrounded by N-terminal ankyrin repeat domains (ARDs), an architecture shared with the canonical transient receptor potential (TRPC) ion channel family. Similarly, structures of the TRP melastatin (TRPM) ion channel family also showed a C-terminal coiled-coil surrounded by N-terminal cytoplasmic domains. This conserved architecture may indicate a common gating mechanism by which modification of cytoplasmic domains can transduce conformational changes to open the ion-conducting pore. We developed an *in vitro* system in which N-terminal ARDs and C-terminal coiled-coil domains can be expressed in bacteria and maintain the ability to interact. We tested three gating regulators: temperature; the polyphosphate compound IP_6_; and the covalent modifier allyl isothiocyanate to determine whether they alter N-and C-terminal interactions. We found that none of the modifiers tested abolished ARD-coiled-coil interactions, though there was a significant reduction at 37°C. We found that coiled-coils tetramerize in a concentration dependent manner, with monomers and trimers observed at lower concentrations. Our system provides a method for examining the mechanism of oligomerization of TRPA1 cytoplasmic domains as well as a system to study the transmission of conformational changes resulting from covalent modification.

## Introduction

The Transient Receptor Potential Ankyrin-1 (TRPA1) ion channel is expressed in nociceptors of the peripheral nervous system [1] where it is activated by a variety of noxious chemical stimuli including electrophilic covalent modifiers [1–3], non-covalent compounds [4], and temperature [5,6]. TRPA1 is also involved in inflammatory signaling [7] and has become an active therapeutic target for treatment of cough [8,9], itch [9,10], and pain [10,11].

Despite the importance of TRPA1 in sensing noxious stimuli, the structural mechanisms of channel activation remain unknown. Since there are multiple channel activators, both covalent and non-covalent, that likely bind to different regions of the channel [3,4,12], it is possible that TRPA1 undergoes different structural rearrangements during activation that depends on the ligand used. Indeed, Cavanaugh, Simkin, and Kim proposed early on that there are different functional states of human TRPA1, one that can be activated by covalent activators in the presence of intracellular polyphosphates and a state that can be activated by Δ^9^-tetra-hydrocannabinol in absence of intracellular polyphosphates but not covalent activators [13]. This suggests the existence of multiple structural states of the channel. Further, it was recently shown using limited proteolysis combined with mass spectrometry that different gating regulators of mouse TRPA1 produced different patterns of proteolysis, consistent with each gating regulator producing unique structural rearrangements [14]. These observations point to the possibility of selectively targeting different activation pathways to regulate the channel. This could prove to be essential for effective pharmacological targeting of TRPA1 where it would be
advantageous to maintain normal sensory function while disrupting pathological pain sensations.

The recently published cryo-electron microscopy structure of human TRPA1 [15] revealed membrane topology of a typical voltage-gated ion channel consisting of six transmembrane domains, where the first four helices make up the voltage-sensing domain (VSD) and the remaining two helices composing the cation selective pore domain (Fig 1). The structure shows no high resolution density for the first ~440 N-terminal amino acids, which contain approximately ten ARDs, as well as portions of the C-terminus [15]. The resolved portion of the cytoplasmic domains consists of a C-terminal tetrameric coiled-coil surrounded by four groups of six N-terminal ankyrin repeat domains (ARDs) (Fig 1), an architecture seen in the TRPC ion channel family structures [16–18], but differing notably from the structure of TRPV1, another TRP channel expressed in nociceptors, that lacks the C-terminal coiled-coil [19–21]. Similar to TRPA1 and TRPC structures, the structures of the TRPM channel family also show a C-terminal coiled-coil surrounded by N-terminal protein domains, though these domains are not ARDs in the TRPM family [22–26].

**Fig 1.**
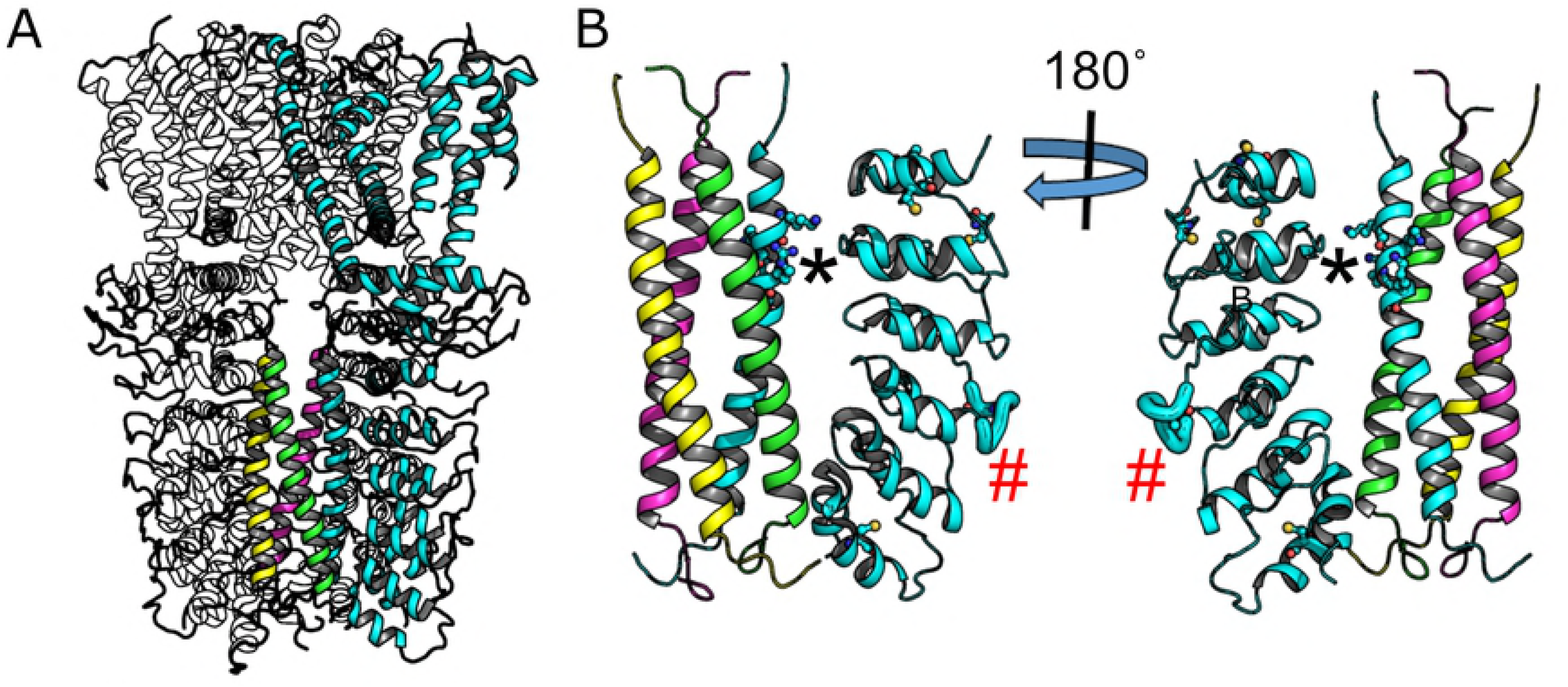
Structural features of human TRPA1. (A) Cartoon structure of human TRPA1 (3J9P) with one subunit highlighted in cyan. The C-terminal coiled-coil helices of all subunits are also shown in color. (B) The cytoplasmic domains showing the resolvable N-terminal ankyrin repeat domains of one subunit along with the coiled-coil. Ball-and-sticks on the coiled-coil helix are positively charged amino acids predicted to destabilize formation of the coiled-coil tetramer. Cysteine residues on the ankryin repeat domains are also shown as ball-and-sticks. The asterisk indicates proposed IP_6_ binding site and the red pound sign indicates the region increased proteolysis upon activation by the electrophilic compound NMM as determined by Samanta et al[14].

Although the TRPA1 cryoEM structure provides a solid starting point to probe structural activation mechanisms, the structure provides no obvious way to determine how conformational changes upon cysteine-modification by electrophilic compounds can be transmitted from the cytoplasmic domains to the ion-conducting pore. Using sequence analysis techniques, Palovcak et al hypothesized the importance of the voltage sensing domain in TRP channel gating by comparing large numbers of sequences of essentially non-voltage gated TRP channels with those of the heavily voltage-dependent K_V_ ion channels [27]. Based on this work, a recent study proposed a common pathway for TRP channel gating through a Ca^2+^ regulated intracellular cavity between the voltage sensor domain (VSD) and the pore domain [28], though no Ca^2+^binding site was observed in the structure. Based on mutation and inter-species chimera approaches, Gupta et al suggest that the S4-S5 linker that bridges the VSD and the pore domain plays an important role in inhibition of human TRPA1 by the synthetic non-covalent channel modifier HC-030031 [4]. Both the Ca^2+^ regulated intracellular cavity and S4-S5 linker are in close physical proximity to the large cytoplasmic domains, and could serve as conduit for conformational changes in the cytoplasmic domains being transmitted to open the ion-conducting pore.

Several studies have shown or implicated that the cytoplasmic domains of TRPA1 are important for regulation by small molecule compounds. A number of cysteine residues in this region have been shown to be the main amino acids involved in activation by irritant compounds such as cinnamaldehyde and allyl isothiocyanate (AITC) [2,29,30]. Recently, it was shown that addition of an electrophilic compound to purified mouse TRPA1 resulted in altered proteolytic accessibility of a loop, highlighted in Fig 1B (red hash mark), between adjacent ARDs [14]. The conformational change required to alter proteolytic accessibility of this loop is unknown. Intracellular polyphosphates are another compound believed to interact with the cytoplasmic domains of TRPA1. It was shown that intracellular polyphosphates were required to maintain channel activation in excised patches [31].

Indeed, it has been reported that IP_6_, a polyphosphate compound, was required for purification of human TRPA1 [15], leading the authors to hypothesized that the negatively charged IP_6_ molecule countered the positively charged amino acids (Fig 1) allowing for tetramerization of the coiled-coil domain [15].

TRPA1 is one of several TRP channels are known to be regulated by temperature [32,33]. The temperature dependence (cold or heat activation) of human TRPA1 remains controversial [5]. Further, whether TRP channels contain a distinct “temperature sensor” or have a diffuse set of amino acids that contribute to differences in heat capacity between the open and closed states [34] remains unknown. However, the temperature dependence of a prokaryotic sodium channel has been shown to be due to unwinding of a C-terminal coiled-coil, leading the authors to suggest a similar mechanism for the coiled-coil of TRPA1 [35]. If this model of TRPA1 temperature sensation is accurate, we should be able to see a temperature dependent unwinding of the TRPA1 coiled-coil. Since IP_6_ is thought to interact with the coiled-coil [15], we might expect that this molecule alters biochemical properties of the cytoplasmic domains at elevated temperatures such as coiled-coil tetramerization, coiled-coil helix stability, or coiled-coil-ARD interactions.

We used isolated protein domains from human TRPA1 consisting of the C-terminal coiled-coil and the N-terminal ARDs to probe the role of IP_6_, temperature, and electrophilic activators on multimerization of the cytoplasmic domains. We showed that coiled-coil concentration is the primary determinant of tetramerization, but with a low affinity such that it is unlikely to be the primary driver of full-length channel tetramerization. We observed that IP_6_ had no effect on the tetramerization of the coiled-coil, suggesting that the requirement of polyphosphates for TRPA1 function in excised patches is not simply due to biochemical stabilization of the coiled-coil. We also showed that the CC helix unwinds ~25% as temperature is elevated, independent of IP_6_, but that the partial helix unwinding had no detectable impact on coiled-coil tetramerization. This is consistent with the model of partial helix unwinding leading to gating as proposed by Arrigoni et al [35]. Finally, we showed that neither removal of IP_6_, increasing temperature, nor addition of AITC abolished interactions between the coiled-coil and the ARDs. The system developed here maintains interactions observed in the full-length channel structure and can serve as a basis in which to study conformational changes in the cytoplasmic domains that result in channel activation.

## Results

### IP_6_ does not alter coiled-coil oligomerization

In order to explore the role of TRPA1 cytoplasmic domains in channel modulation, we developed constructs suitable for biochemical characterization. The primary sequence of the human TRPA1 coiled-coiled consisting of amino acids A1036-T1078 (referred to as CC1) contains no tryptophan residues and few other residues that absorb at 280 nm making it difficult to observe in standard size exclusion chromatography with absorbance detection. Hence, in order to examine coiled-coil oligomerization we expressed CC1 as a maltose-binding protein (MBP) fusion (referred to as MBPCC1, Fig 2A). In addition to providing strong absorption signal at 280 nm this also allows for easy discrimination between monomeric fusion protein of ~50 kDa and tetrameric protein of ~200 kDa using size exclusion chromatography (SEC). MBPCC1 expressed robustly in E. coli and was used as a means to monitor oligomerization of the coiled-coil (Fig 2).

**Fig 2.**
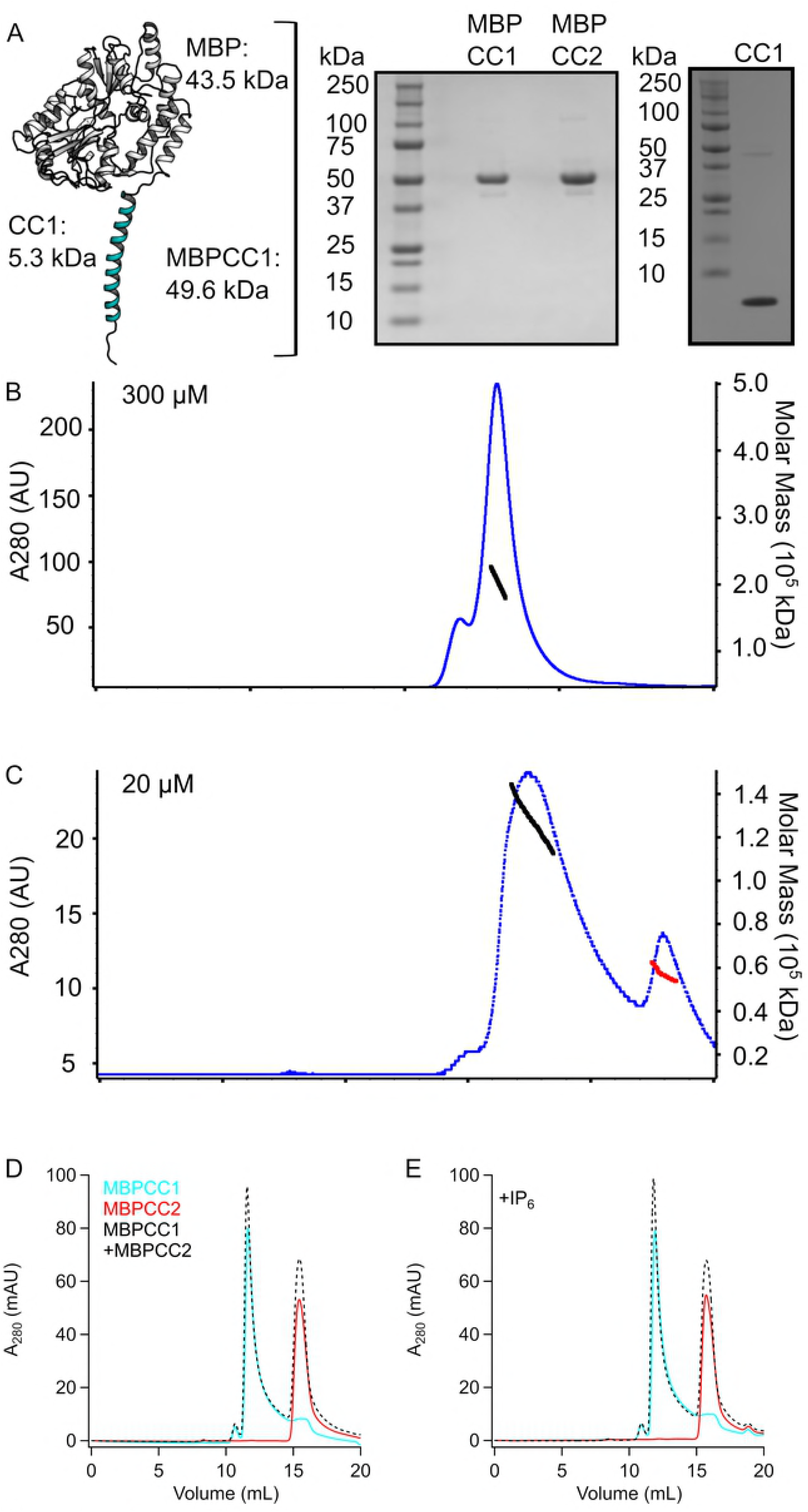
Oligomerization of hTRPA1 coiled-coil. (A) Cartoon depicting the MBPCC1 fusion protein along with Coomassie-stained gels of purified MBPCC1, MBPCC2, and isolated CC1 protein. (B) SEC-MALS showing high concentration of MBPCC1 forming a tetramer (Mw calculated to be 203 kDa +/- 1.5%) in the absence of IP6. (C) SEC-MALS showing MBPCC1 at lower concentration is no longer tetrameric but containing a mix of trimers (Mw calculated to be 136 kDa) and monomers (Mw calculated to be 53 kDa) in the absence of IP_6_. For panels in (B) and (C), blue dots indicate absorbance at A280 and black or red dots indicate regions and values where Mw was calculated. (D), (E). SEC chromatograms of MBPCC1, MBPCC2, and a mixture of MBPCC1+MBPCC2 in the absence (D) or presence (E) of IP6 showing no clear signs of interaction between CC1 and CC2.

We used size-exclusion chromatography combined with multi-angle light scattering (SEC-MALS) to get an accurate determination of the molecular weight of MBPCC1. MBPCC1 protein at 15 mg/ml (~300 μM) in the absence of IP_6_ ran at a molecular weight consistent with a tetrameric protein on an SEC column and the molecular weight determined from light scattering was 203 kDa (+/- 1.5%) (Fig 2B). Tetrameric protein was observed in the absence of IP_6_ during all stages of expression and purification, indicating that IP_6_ was not required for coiled-coil solubility or tetramerization under our experimental conditions. When we diluted MBPCC1 to 1 mg/ul (~20 μM) and analyzed the protein with SEC-MALS we observed protein at 138 kDA (± 0.4%), a molecular weight in between that of a dimer and trimer as well as a peak consistent with a monomer (53 kDa) (Fig 2C). Thus, multimerization appeared to depend on the concentration of protein but not IP_6_.

When we ran the human TRPA1 protein sequence through the COILS algorithm[38], we noticed a second region consisting of amino acids D1082-K1113 (referred to as CC2) that showed propensity for forming a coiled-coil. We tested whether a purified protein fragment corresponding to this region formed a coiled-coil *in vitro* by expressing it as an MBP fusion protein (MBPCC2). When MBPCC2 was run on SEC in the absence or presence of IP_6_ it eluted at a molecular weight consistent with a monomer (Fig 2D,E). When we incubated purified MBPCC1 at a low concentration and MBPCC2 together and ran the sample on SEC, the chromatograms showed no sign of heteromeric oligomerization between CC1 and CC2 in the absence or presence of IP_6_ (Fig 2D,E).

### Isolated coiled-coil protein is helical and unwinds at room temperature

It was recently shown that the unwinding of a C-terminal coiled-coil at increasing temperatures underlies temperature-sensitive gating of a prokaryotic sodium channel and it was suggested that a similar mechanism could be the case for TRPA1 [35]. We tested whether temperature would partially unwind CC1 and whether IP_6_ would prevent this unwinding, testing the hypothesis that the functional requirement for IP_6_ in excised patches is due to its stabilization of CC1.

We used circular dichroism spectroscopy (CD) on isolated CC1 (Fig 2A) to probe the helical content at increasing temperatures in the presence and absence of IP_6_ (Fig 3). From 4°C to 47°C, there was a marked and reversible decrease in ellipticity of CC1 in the presence (Fig 3A) and absence of IP_6_ (Fig 3B). When the ellipticity at 222 nm at different temperatures was normalized to the ellipticity at 222nm at 4°C we observed a reversible ~25% reduction, as temperature is increased to 42°C indicating that part of the coiled coil was reversibly lost as temperature was increased and that this occurred in an IP_6_-independent manner. Although these data are not sufficient to conclude that the partial unwinding of CC1 contributes to the temperature-dependent gating of TRPA1, they are consistent with the model proposed by Arrigoni et al [35].

**Fig 3.**
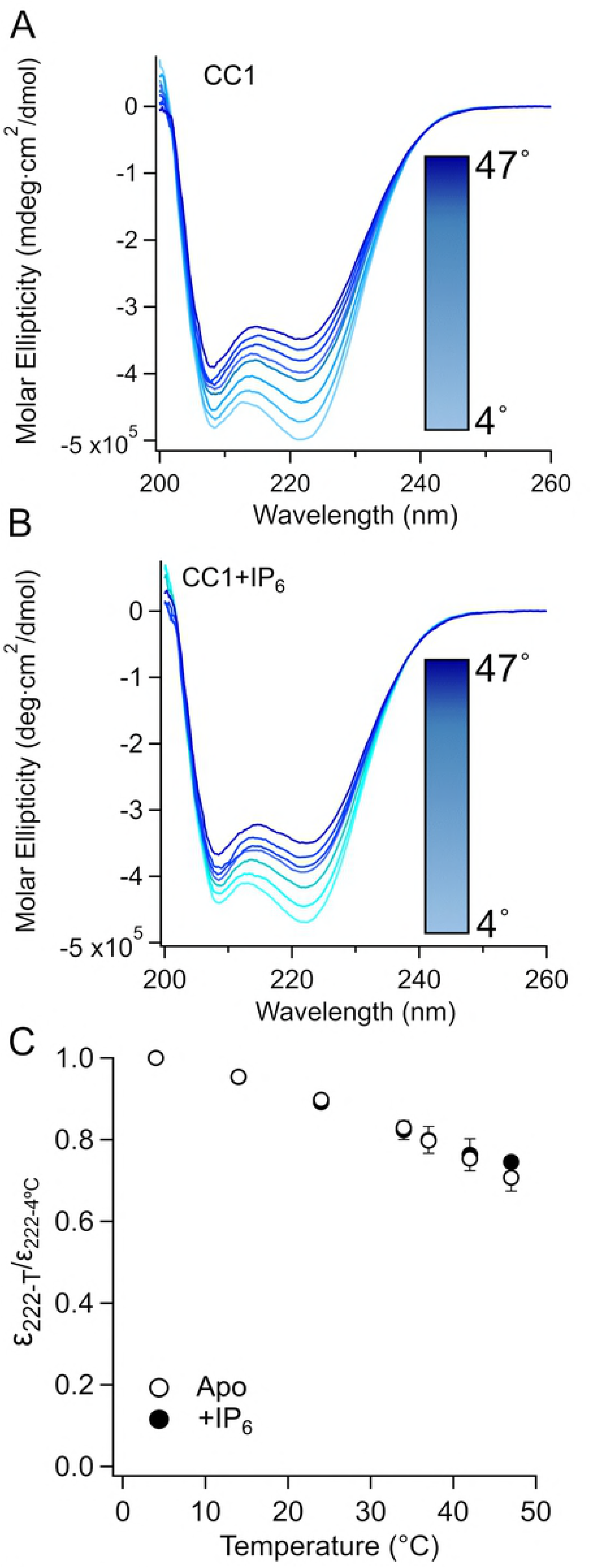
Helical unwinding of isolated CC1. (A and B) Far-UV CD spectrum of CC1 at different temperatures (4°C, 14°C, 24°C, 34°C, 37°C, 42°C, 47°C) in the absence (A) or presence (B) of IP_6_. C. Normalized ellipticity (to ellipticity measured at 4°C) at 222 nm (n=2).

### Concentration but not temperature or IP_6_ determines CC1 multimerization

We next sought to determine whether temperature would have an effect on oligomerization of CC1 in the presence and absence of IP_6_. MBPCC1 fusion protein without IP_6_ was incubated at 4°C, 24°C, and 32°C and run on an SEC column equilibrated at the same temperature. These temperatures were chosen in part due to the amount of helical unwinding noticed in CD experiments as well as instrument limitations at higher temperature. Neither changing temperature nor IP_6_ altered tetramerization as determined from the SEC profiles (Fig 4A), demonstrating that neither temperature nor IP_6_ was a major factor in coiled-coil tetramerization under our conditions.

**Fig 4.**
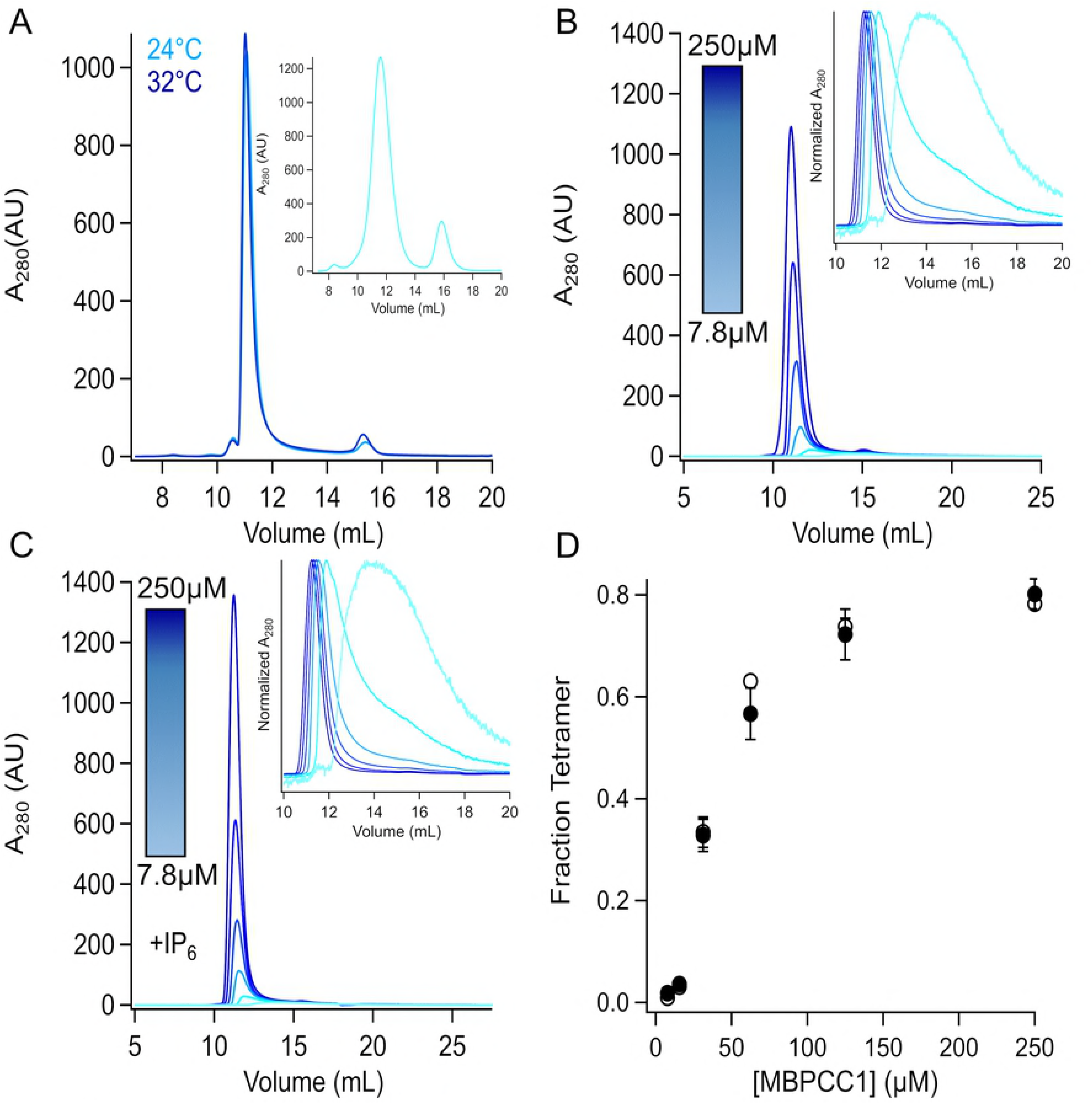
Concentration, not temperature or IP_6_, is the primary driver of tetramerization. A. SEC chromatograms of MBPCC1 at 4°, 24°, and 32°C. The instrument and column for T=24°C and T=32°C were the same (Superdex 200 Increases run on a Shimadzu system), but different from sample injected at 4C (Superdex 200 column and Akta Explorer). Both peaks are at molecular weights consistent with tetrameric protein at all temperatures tested. B and C. Protein at different concentrations (250 μM, 125 μM, 62.5 μM, 31.25 μM, 15.625 μM, and 7.8125 μM) was injected at 24°C in the absence (B) and presence (C) of IP_6_. Insets show the normalized absorbance to highlight the shift in elution volume at lower concentrations. (D) Fraction of tetramer vs concentration (n=3 for each concentration).

The TRPA1 coiled-coil forms intersubunit interactions (Fig 1) that may contribute to the tetramerization of the full-length channel. It has previously been shown that the intracellular T1 domains of some voltage-gated potassium channels specify compatibility for tetramerization among different K_V_ subunits [39,40]. We therefore tested whether the TRPA1 coiled-coils could drive full-length channel tetramerization. We evaluated tetramerization of decreasing concentrations of MBPCC1 fusion protein in the presence or absence of IP_6_ using size-exclusion chromatography (Fig 4C, D). There fraction of tetrameric MBPCC1 at different concentrations was the same in the presence or absence of IP_6_ (Fig 4E). The concentration-tetramerization curve was half maximal in the concentration range of tens of micromolar. It seems unlikely that the micromolar concentrations required for tetramerization are the driver of full-length channel tetramerization. Rather, the tetramerization of the transmembrane domain likely increases the local concentration of CC1 to induce tetramerization of coiled-coil.

### Temperature, IP_6_, electrophilic compounds do not alter ARD-CC1 binding

Since the full-length TRPA1 structure showed interactions between the C-terminal coiled-coil and the N-terminal ARDs we tested if our isolated coiled-coil protein could interact with isolated ARDs *in vitro.* To test this, we made a His-tagged construct containing amino acids 446-639, corresponding to ARDs with resolvable density in the human TRPA1 cryoEM structure. We then co-expressed this ARD construct with a His-tagged MBPCC1 construct or expressed ARD alone and tested whether the proteins copurified with amylose affinity resin (Fig 5). As shown in the Western Blot analysis in Fig 5, we observed that ARD protein could be co-purified with MBPCC1 using amylose affinity chromatography with little or no ARD protein bound to amylose resin in absence of MBPCC1 (Fig 5).

**Fig 5.**
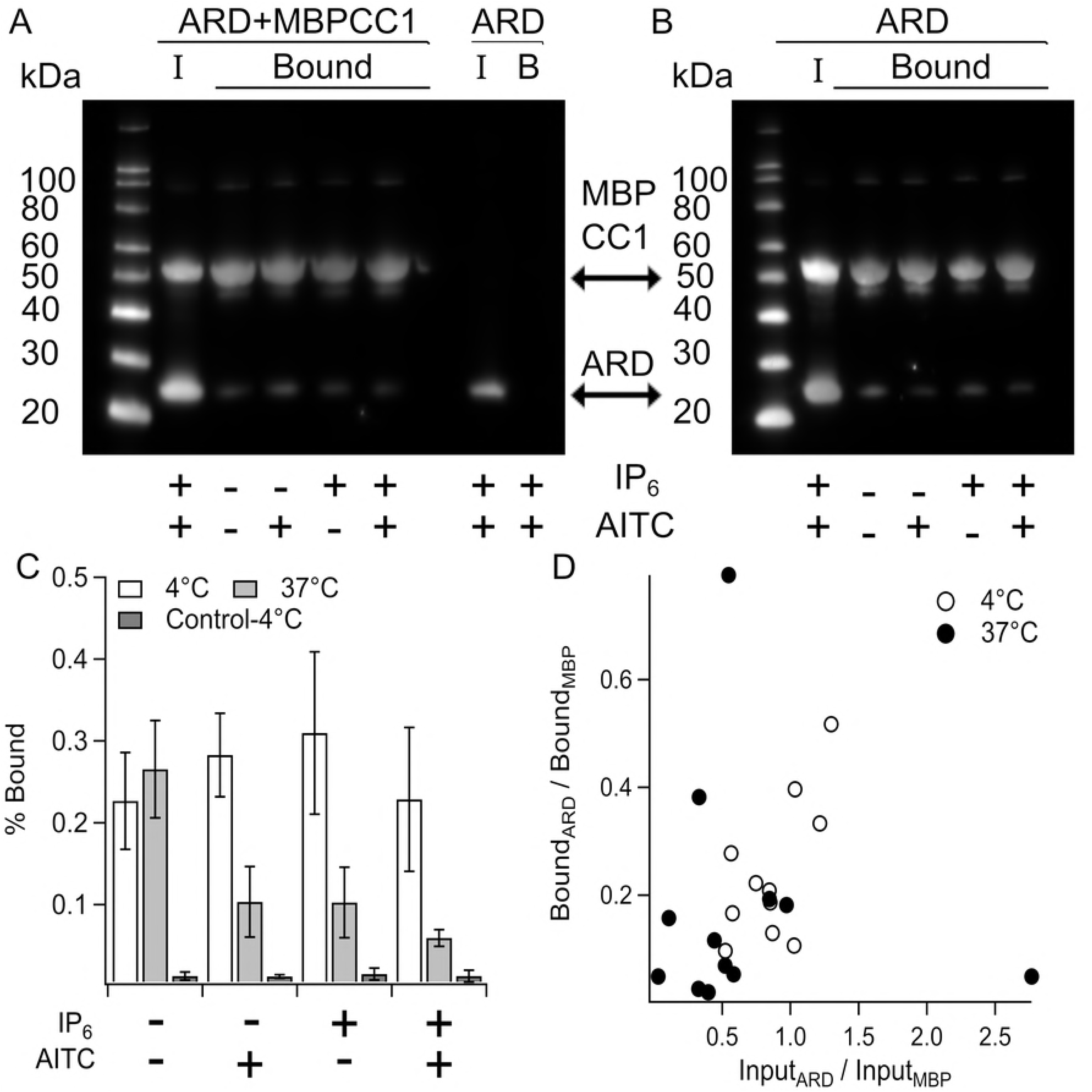
Interaction of MBPCC1 with the N-terminal ARD. A and B. Anti-His Western blots of amylose-resin purified protein from bacteria expressing both ARD and MBPCC1 or ARD alone. Incubation with resin and subsequent washing of resin was carried out at 4°C (A) or 37°C (B). “I” indicates input sample and “Bound” indicates protein pulled down that remained after 4 wash steps. At each temperature lysates were incubated with IP6 and/or AITC. C. Fraction of ARD bound to MBPCC1 normalized to input (see methods). Control indicates amylose purification of cells only expressing ARD at 4°C. D. Bound ARD/MBP ratio vs Input ARD/MBP ratio showing that more ARD in the input correlates with increased ARD binding to MBPCC1 during purification.

The TRPA1 cryoEM structure contains a non-protein density at the interface between the coiled-coil and ARDs that was attributed to IP_6_ (Fig 1B). We therefore tested whether IP_6_, temperature, and the electrophilic TRPA1 activator AITC altered MBPCC1-ARD interactions (Fig 5). Although AITC was added in the presence of intracellular proteins (i.e. cleared cell lysates) and reducing agent (2 mM TCEP), modification of cysteines by electrophilic activators occurs even in the reducing environment in intact cells/tissue *in vivo* to produce noxious sensation. At 4°C, there was no difference in the amount of ARD that co-purified with MBPCC1 in the absence or presence of IP_6_ and AITC (Fig 5A,C). We next tested whether increasing temperature to 37°C would change binding (Fig 5B,C). Although there was a significant decrease in the amount of ARD pulled-down with CC1at 37°C, binding was nonetheless above background levels (Fig 5B,C). When we plotted the ratio of ARD to MBPCC1 of input sample to the ratio of ARD to MBPCC1 in the amylose-bound sample we observed a correlation between the ratio of ARD:MBPCC1 expressed and the fraction of ARD pulled-down. Notably, the ARD expression used in 37°C experiments was generally less than that of experiments at 4°C (Fig 5D), suggesting a possible explanation for the lower co-purification we observed. In any case, at both 4°C (Fig 5A, C) and 37°C (Fig 5B, C) the ability of the ARDs to interact MBPCC1 was maintained in the presence or absence of IP_6_ or AITC.

## Discussion

In this study we aimed to develop a reduced *in vitro* system in which we could reconstitute the interactions of the N-terminal ARDs and the C-terminal coiled-coil observed in the full length human TRPA1 structure [15]. Our results show that the C-terminal coiled-coil can form tetramers (Fig 2 and Fig 4) and interact with the N-terminal ARDs (Fig 5) as observed in the full-length channel structure, key requirements for our *in vitro* system.

We used our system to test a number of hypotheses that could provide insight into how the cytoplasmic domains could be involved in channel gating. We showed that IP_6_ was not required for structural stability of the TRPA1 C-terminal coiled-coil (Fig 2, 4) or the oligomeric stability of the interactions between the N-terminal ARDs and the C-terminal coiled-coil (Fig 5). We further showed that AITC was not required for the interaction between the ARDs and the coiled-coil in our isolated-domain system (Fig 5). We showed a reversible partial coiled-coil helix unwinding as temperature was increased (Fig 3), consistent with a model proposed by Arrigoni et al [35], but this helix unwinding did not fully abolish coiled-coil-ARD interactions (Fig 5). Helix unwinding may result in different physical space being occupied, pushing away or bringing closer the N-terminal ARDs. This can be accommodated by flexibility in the ARDs such as that observed in mouse TRPA1 where an electrophilic activator altered the proteolytic accessibility of a loop between ARDs [14]. However, it is important to note that we cannot determine if this helix unwinding is involved in channel gating from our data. Together these data suggest that the role of intracellular polyphosphates and cysteine-modifying electrophilic compounds are more complex than serving as just stabilizing ligands for N-terminal and C-terminal domain interactions.

## Materials and Methods

### Molecular biology

Human TRPA1 cDNA was a gift from Ajay Dhaka. CC1 (amino acids A1036-T1078) and CC2 (amino acids D1082-K113) were cloned into the Nco I and Hind III restriction sites in the pHMAL-c2TEV vector which contains an N-terminal poly-histidine tag followed by maltose-binding protein (gift from WNZ). The TRPA1 ARD construct (amino acids 446-639) was cloned into the pET28b vector using Nhe I and Sac I restriction sites in frame with an N-terminal poly-histidine tag.

### Fusion protein expression and purification

MBP fusion proteins in the pHMAL-c2TEV vector were transformed into BL21(DE3) competent cells and grown at 37°C to OD_600_ between 0.5-0.75 and protein expression was induced with 0.5 mM IPTG for 3.5 hours at 37°C. Harvested cells were suspended in 50 ml/L culture Buffer A1T (150 NaCl, 20 TrisHCl, pH 7.8, 2 mM TCEP) and stored at −20C until needed. Thawed cells were lysed via sonication after a ten minute incubation with a protease inhibitor cocktail containing PMSF (1 mM), Aprotinin (1 μg/ml), pepstatin (3 μg/ml), leupeptin (1 μg/ml). Lysed cells were cleared via centrifugation for 35 minutes at 30,000 x *g* in a Beckman JA-20 rotor. Cleared lysates were incubated with amylose resin (NEB) for 60 minutes at 4°C and purified using gravity flow. Resin was washed with at least 15 column volumes of Buffer A1T and eluted with Buffer A1T supplemented with 20 mM maltose (sigma). Protein was concentrated and subjected to size exclusion chromatography to remove maltose and used promptly for assays or stored at −20°C for future use.

Analytical size exclusion chromatography was performed using a Shimadzu HPLC with a Superdex 200 Increase column with Buffer A1T at room temperature (24°C). For SEC experiments at 32°C, protein was incubated at 32°C for 60’ and run on SEC using column oven set to 32°C. Large scale preparative SEC runs carried out at 4°C showed no clear difference between those at 24°C though extensive analysis was not carried out at this condition. To determine the fraction of tetramer at different protein concentrations, the area under the absorption curve corresponding to the volume range where a gaussian fits at high concentration is divided by the total of area of protein absorption.

To isolate coiled-coil protein, TEV protease was added to protein at 1:500 dilution and incubated two hours at room temperature. The digested protein was then run over a HisPur cobalt column to remove excess MBP protein and dialyzed into a buffer containing 100 mM NaCl, 20 mM Tris-HCl, pH 7.4 with or without IP_6_.

### Multi-Angle Light scattering

Size-exclusion chromatography (Superdex 200 column) coupled with light scattering, refractive index, and ultraviolet absorption (SEC-LS-RI-UV) was done under the SEC-MAL system, which consisted of a P900 HPLC pump (GE), a UV-2077 detector (Jasco), a Tri Star Mini Dawn light scattering instrument (Wyatt), and an Opti Lab T-Rex refractive index instrument (Wyatt).

### Circular dichroism spectroscopy

CD spectra of CC1 were collected on a Jasco J-1500 CD spectrometer. Samples contained 0.1 mg/ml (~20 μM) protein in buffer containing 100 mM NaCl, 20 mM Tris-HCl, pH 7.4 with or without IP_6_. Samples were equilibrated at the indicated temperature for at least 10 minutes before measurement and measured in a 0.1-cm cuvette. Measurements were taken in continuous scanning mode with a scanning rate of 50 nm/min, data integration time of 2 s, and bandwidth of 1 nm. Data presented are an average of three scans, and data at wavelengths resulting in a high-tension value at or above the recommended 800-V cutoff were excluded.

### Binding Assays

For binding assays ARD or ARD+MBP-CC1 was transformed into BL21(DE3) competent cells and grown at 37°C to OD600 ~0.75. Protein expression was induced with .75 mM IPTG and cells were transferred to 25°C for 20-24 hours. Harvested cells were suspended in Buffer A1T and stored at −20°C until used for purification. Thawed cells were lysed via sonication after a ten-minute incubation with a protease inhibitor cocktail containing PMSF (1 mM), Aprotinin (1 μg/ml), pepstatin (3 μg/ml), leupeptin (1 μg/ml). Lysed cells were cleared via centrifugation for 35 minutes at 30,000 x *g* in a Beckman JA-20 rotor. Cleared lysates were added to 50 μL of equilibrated amylose resin and incubated at either 4°C or 37°C for 60 minutes. Resin was washed four times with 500 μL Buffer A1T and eluted with Buffer A1T supplemented with 20 mM maltose. For binding assays in the presence of IP_6_ and/or AITC, each compound was added at least 10 minutes prior to lysis and included in both wash and elution buffers.

Since both MBP-CC1 and ARD contained N-terminal His-tags, they were both probed on Western Blot with anti-His primary antibody (QIAgen) overnight at room temperature in TBST supplemented with 5% milk. Membranes were washed three times for five minutes with TBST and incubated with HRP conjucated anti-mouse IgG secondary antibody for 60’ at room temperature. Membranes were then washed three times for five minutes with TBST and imaged after addition of ECL femto reagent. For quantification, ImageJ was used to determine regions of interest around each protein band. To determine the fraction of ARD bound, we took the ratio of amylose-purified ARD signal to input ARD signal and normalized by the fraction of MBP bound compared to input to compensate for having excess protein in the cell lysates. Each Western Blot used for the analysis of fraction bound ARD contained both input and amylose purified samples on the same blot. Thus, the Blots shown in Fig 5 were primarily used for illustrative purposes (though each contained one sample that could be used for quantification).

## Acknowledgments

We would like to thank members of the Gordon and Zagotta labs for comments and discussions throughout the project. We thank Ajay Dhaka for the human TRPA1 DNA.

